# Living apart if you can – how genetically and developmentally controlled sex has shaped the evolution of liverworts

**DOI:** 10.1101/2021.02.08.430207

**Authors:** Xiaolan He, Jorge R. Flores, Yu Sun, John L. Bowman

**Affiliations:** Finnish Museum of Natural History, University of Helsinki, FI-00014, Helsinki, Finland; Lushan Botanical Garden, Chinese Academy of Sciences, Jiujiang, Jiangxi 332900, China; School of Biological Science, Monash University, Melbourne VIC 3800, Australia

**Keywords:** bryophytes, cytology, evolution, genomics, haploid dioicy, liverwort, phylogeny, sex chromosome, sex-determination, sexual system

## Abstract

Sexual differentiation in bryophytes occurs in the dominant gametophytic generation. Over half of bryophytes are dioicous, and this pattern in liverworts is even more profound as over 70% of species are dioicous. However, the evolutionary mechanisms leading to the prevalence of dioicy and the shifts of sexual systems between dioicy and monoicy have remained poorly known. These essential factors in reproductive biology are explored here in light of phylogenetics combined with evidence of genomic characterization of sex chromosomes and sex-determination, as well as cytology. Our analyses and discussions on liverworts are focused on: (1) ancestry and shifts in sexuality, (2) evolution of sex chromosomes and maintenance of haploid dioicy, and (3) environmental impact on the evolution of monoicism. We show that the dioicous condition is ancestral in liverworts, and the evolution of sexual systems is both conserved and stable with an ancient origin, but also highly dynamic in certain more recently diverged lineages. We assume that the haploid dioicy maintained in the course of evolution must be linked to the genetically controlled sex-determination, and transition from genetically to developmentally controlled sex determination, the evolution of monoicism, is driven by ephemeral and unstable environments. Monoicy is less stable in the long-term than dioicy, and thus, ultimately, dioicy is selected in liverworts. It is concluded that sexual dimorphism is maintained through a highly dynamic evolutionary process, sex chromosomes bearing distinct set of evolutionary forces that can have large impacts on genome evolution and may also promote speciation.

## Introduction

Sexual reproduction and dispersal are two interacting features playing an essential role in the evolution of almost all eukaryotes. This is particularly true for bryophytes due to their unique life cycle characteristics and sexual systems. The life cycle of bryophytes is distinct from vascular plants; in bryophytes the haploid phase consists of a free-living gametophyte responsible for both sexual and asexual functions, and the diploid phase represented by a sporophyte that produces large amounts of haploid resistant spores as dispersal units. Although the sexual stage occurs only in the gametophytes, in terms of distance, gene flow achieved through spore dispersal is orders of magnitude greater than that through sperm dispersal (Wyatt, 1982). Therefore, production of sporophytes has widely been considered as a key factor for measuring success in bryophyte reproduction. Sexual reproduction in bryophytes can be accomplished by either dioicous or monoicous species and infrequently in some species both reproductive modes occur. The sexual condition strongly affects production of sporophytes, with dioicous species producing sporophytes less often than monoicous species, and a fair number of dioicous moss and liverwort species have never been found to produce sporophytes (Schuster, 1966; Longton, 1976).

More than half of the bryophytes are dioicous, and this pattern in liverworts is even more profound as dioicy is represented by over 70% of species (Schuster, 1966; Longton, 1976). The only exception is the hornworts, among which 60% of species are monoicous (Villarreal & Renner, 2013). The evolutionary mechanisms leading to the prevalence of dioicy in liverworts, and in mosses as well, remain unexplained, but it is believed that outcrossing in promoting and maintaining genetic variability should be selected in bryophytes, as dioicy simply makes outcrossing obligatory (Mishler, 1988; Longton, 2006). However, intra-gametophytic self-fertilization in monoicous bryophytes leading to a totally homozygous sporophyte in one generation, will result no or less inbreeding depression in future generations because of efficient purging of deleterious alleles (Eppley *et al*., 2007; Taylor *et al.*, 2007; Szövényi *et al*., 2014). Sexual reproduction in bryophytes has also been considered facultative with genetic variation maintained largely through somatic mutation, because asexual reproduction is effective in rapid spreading of existing populations and production of sporophytes is infrequent in many dioicous species (Mishler, 1988). In contrast, the study of García-Ramos *et al*. (2007) emphasizes the role of asexual reproduction in promoting coexistence of the sexes. For diploid and polyploid organisms, theory predicts that large population with stable reproductive systems can be highly variable even with only small number of sexually reproducing individuals per generation (Bengtsson, 2003). Nonetheless, both empirical and theoretical studies indicate that the key advantage of sexually produced offspring is the ability to colonize unpredictably available new habitats, which is impossible to achieve through asexual reproduction (Longton, 2006; García-Ramos *et al.*, 2007).

The genetic basis for dioicism of bryophytes, and the role of sex chromosomes, has been debated for the past century (Lewis, 1961; Smith, 1978; Ramsay & Berrie, 1982). The correlation between dimorphic chromosomes in size and phenotypic sexual expression found in the liverwort genus *Sphaerocarpos* by Charles E. Allen (1917, 1919) was the first direct evidence of the occurrence of sex chromosomes in bryophytes. Other presumed sex chromosomes widely reported in bryophytes in subsequent investigations were mostly based on their presence in one or the other sex and the amount and distribution of heterochromatin, the nuclear material that remains highly condensed within the interphase nucleus (Heitz, 1928). He named the large heterochromatic sex chromosomes of dioicous *Pellia neesiana* (n = 8 + X/Y) as macrochromosomes, whereas microchromosomes are for the small heterochromatic sex chromosomes of dioicous *P. endiviifolia* (n = 8 +x/y). Tatuno (1941) renamed the macrochromosomes of Heitz as ‘H’, and the microchromosomes as ‘h’. Subsequently, the available cytological evidence suggested that sex chromosomes tend to bear high content of heterochromatin, as in *Sphaerocarpos* and also other animal species (Heitz, 1928, 1933; Lorbeer, 1934; Tatuno, 1933; Segawa, 1965; Newton, 1977). However, Tatuno (1941), who assumed the monoicous condition is primitive in liverworts, noted both H- and h-chromosomes in some monoicous species, indicating that not all H or h chromosomes are sex chromosomes.

Mutagenesis experiments whereby female plants could be transformed into male plants, but not vice versa provided evidence that there exists a ‘feminizer’ locus on the female sex-specific chromosome in *Sphaerocarpos*, *Marchantia* and *Pellia* (reviewed in Bowman, 2016; Heitz, 1949). Likewise, plants possessing a haploid autosomal complement along with both sex chromosomes (A+XY) were functionally female, indicating that the feminizer locus is dominant. Further, as the transformed males (possessing a X chromosome) had immotile sperm suggested that there exist multiple sperm ‘motility’ loci on the male sex-specific chromosome of *Sphaerocarpos* and *Marchantia* (reviewed in Bowman, 2016; Heitz, 1949). These are the most convincing data on the sex-determining role of specific chromosomes in liverworts. Following these insights, a stasis of several decades ensued until recently, after the sequences of sex chromosomes of a liverwort *Marchantia polymorpha* (Yamato et al 2007; Bowman *et al*., 2017) and a moss *Ceratodon purpureus* (McDaniel *et al*., 2013a; Carey *et al*., 2020) were characterized.

Whether dioicy is the ancestral condition of bryophytes and how genetically controlled sex-determination affects the evolution of sexual systems await more investigations. Results from early cytological and cytogenetic studies were largely constrained by uncertainty on the direction of evolution of the sexual systems and on the ploidy level of liverworts (Berrie, 1960; Newton, 1984). To explain these essential questions in reproductive biology, historical approaches combined with systematics must be employed. The former question has been widely discussed and long debated, and either dioicy or monoicy has been assumed to be ancestral for bryophytes (Smith, 1978; Anderson, 1980; Wyatt & Anderson 1984; Newton, 1983, 1986). Limited evidence derived from phylogenetic analyses has given different results for mosses and liverworts, respectively. McDaniel *et al.* (2013b) proposed a high lability of sexual systems in mosses, whereas Laenen *et al.* (2016) suggested dioicy as the putative ancestral state of liverworts, yet the inferred phylogenetic signal associated with the sexual system is significant only at the deepest nodes. Because the evolution of sexual systems may have proceeded differently in different lineages, any analysis attempting for the overall pattern of a group may not be rigorous enough to reveal the true or hidden revolutionary process, therefore different lineages within the group should also be counted separately. The second question on the impact of genetically controlled sex-determination on the evolution of sexual systems in bryophytes has not been explored previously, obviously due to limitation of available data that could be used to address questions as such. At present, despite that genome level evidence on sex chromosomes of bryophytes is available only from two species, it is sufficient to use it and also other evidence to analyze and discuss further on the potential evolutionary mechanisms and environmental influence responsible for the resultant pattern of sexual systems, and to identify profitable ways of further research. Here we will focus our analyses and discussions on liverworts: (1) ancestry and shifts in sexuality in a phylogenetic context, (2) evolution of sex chromosomes and maintenance of haploid dioicy, and (3) environmental impacts on the evolution of monoicism. Directions for future research are proposed.

### The ancestry, and highly conserved and stable dioicy

Liverworts encompass over 7000 species in nearly 400 genera and 90 families (Söderström *et al*., 2016). Phenotypic variations of the gametophytes of liverworts are the most prominent among bryophytes, represented by three highly distinctive types of body plan referred to as simple thalloid, complex thalloid and leafy organizations, coupled with numerous unique structures. The life span of a gametophyte varies from ephemeral to perennial, and the maturation of a sporophyte can last from a few weeks to nearly one year. The sporophytes present less extensive morphological variation than those of the gametophytes, but their meiotic pattern in sporogenesis is extremely varied (Brown & Lemmon, 2013). Liverworts occupy a vast geographical range in all terrestrial environments on substrates from bare soil on dry land to tall canopies as epiphytes of rainforest. Although they share with other bryophytes the same life cycle characteristics, liverworts differ markedly from mosses in possessing a low number and narrow range in chromosome numbers, with over 85% of species having the basic number n=8, 9 or 10 (Berrie, 1960; Newton, 1983, 1988), and with only a rare occurrence of polyploidy (ca. 5% of the species, Laenen *et al*., 2016).

To further understand the evolutionary mechanisms leading to the predominant dioicy, and shifts between sexual systems in liverworts, we performed phylogenetic analyses to reconstruct the evolution of sexual system through time across 80% of the extant liverwort genera and 97% of families (Figs. 1–3). Each major clade was also analyzed separately for detection of any specific pattern (Figs. 2 and 3). In our phylogeny, dioicy is ancestral in liverworts and it has persisted as the dominant condition throughout their evolutionary history (Marchantiphyta, Fig. 3). The same pattern is seen in Pelliidae, which includes the simple thalloid liverworts and a few leafy species, and in Jungermanniales, one of the major leafy liverwort groups. The most frequent shifts of the sexual systems occurred from dioicy to monoicy (e.g., in Marchantiopsida during the Cretaceous; Fig. 3), followed by reversals from monoicy to dioicy (e.g., in Porellales during the Quaternary; Fig. 3). Shifts from dioicy to monoicy were infrequent in Pelliidae and Jungermanniales and occurred mostly in the recently diverged nodes. These monoicous species are associated with polyploidy, such as in the genera *Chiloscyphus, Calypogeia* and *Nardia* in the leafy liverworts, and in *Metzgeria* and *Pellia* in the simple thalloid liverworts. We conclude that the shift to monoicy in these two major liverwort clades resulted from autopolyploidy or allopolyploidy, possibly due to somatic doubling involving a failure of mitosis that produces diploid cells in the gametophytes or tetraploid cells in the sporophytes, or diplospory involving a failure of meiosis in the sporophyte (Wyatt & Anderson, 1984). Polyploidy evolved from apospory, commonly presented in mosses is unlikely to happen in liverworts (Smith, 1978; Anderson, 1980), as induced apospory has only been reported for *Blasia pusilla* (Raudzens & Matzke, 1968).

**Fig. 1.**
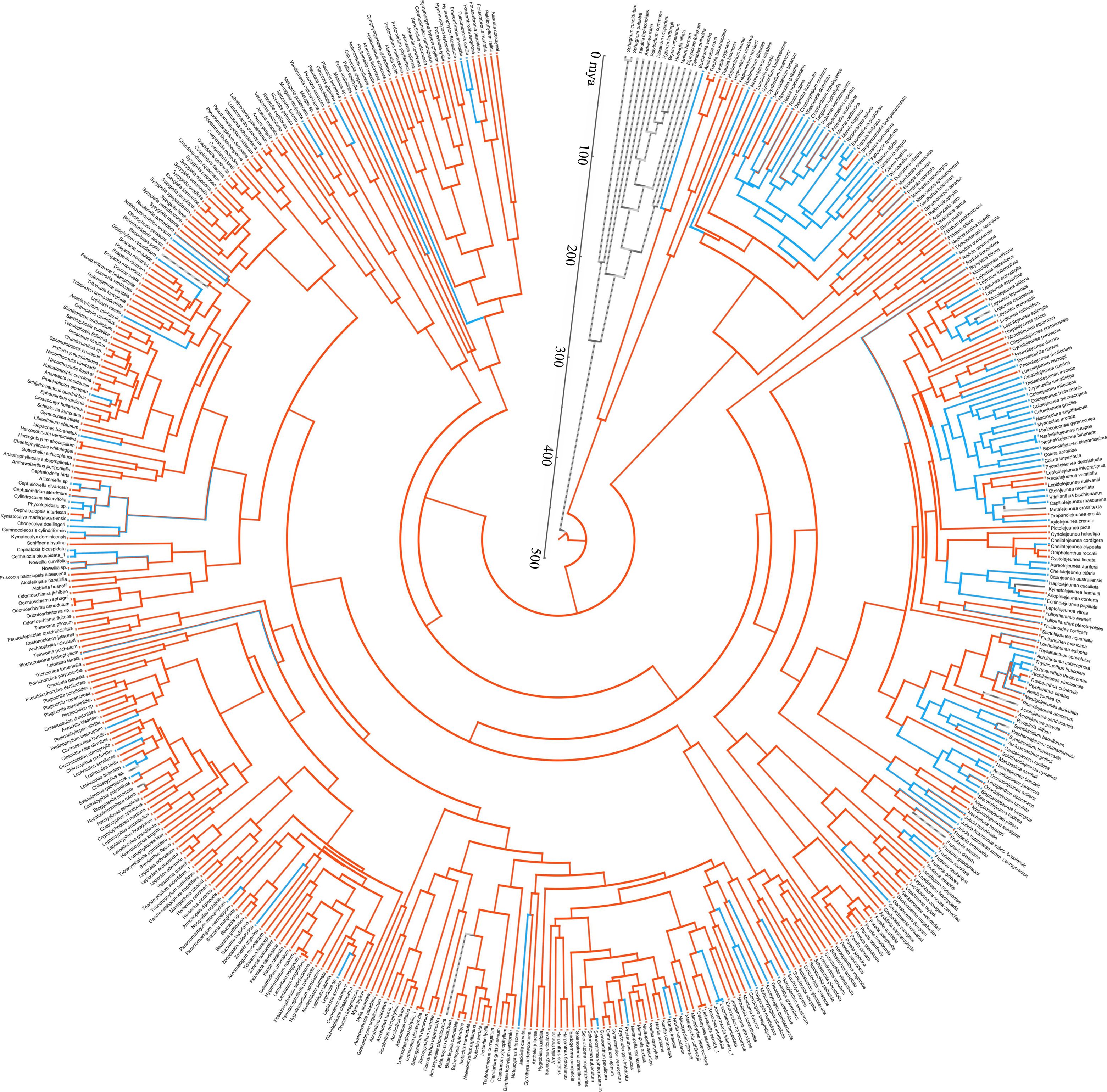
Evolutionary pattern of sexual systems of liverworts. The dated phylogenetic reconstruction was performed using 466 species representing 299 liverwort genera and 84 families based on Bayesian Inference, as implemented in BEAST v1.10.4 (Suchard *et al*., 2018). Of 3232 aligned bases of nucleotides represent three markers (*rbc*L, *rps*4 and *trn*L-F). The taxon sampling represents 80% of the extant liverwort genera and 97% of the families (Söderström et al., 2016). The ancestral character state condition for the sexual system was reconstructed throughout the phylogeny on the basis of maximum parsimony in MESQUITE 3.6.1 (Maddison & Maddison, 2019). The evolution of the sexual system through time was shown as dioicous (orange) and monoicous (blue). Detailed information on the material and methods and the dated phylogeny can be found in the Supplementary Material as S1 and Fig. S1 respectively.

**Fig. 2.**
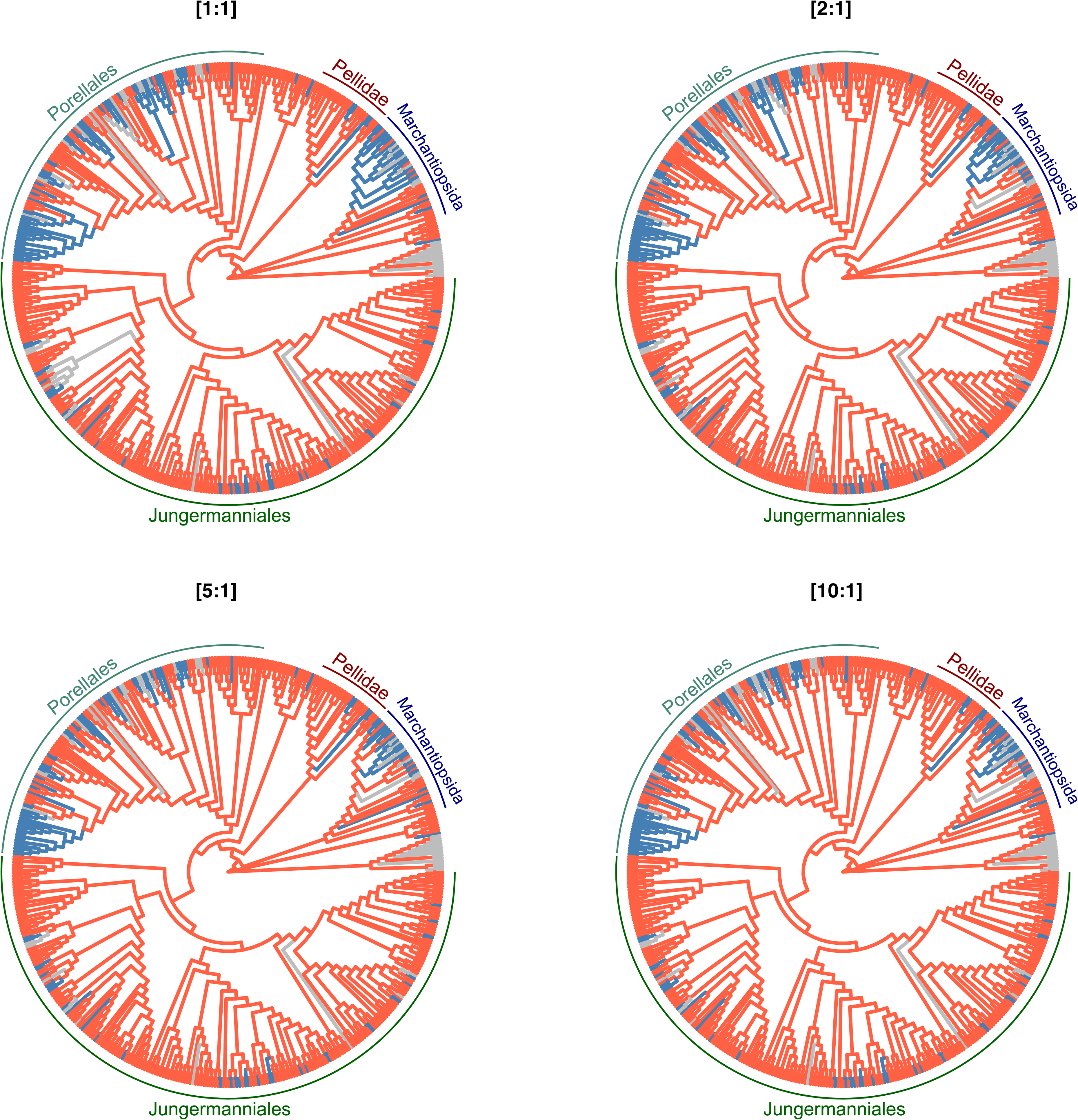
Evolution of sexual systems of liverworts under different scenarios. Reversal costs were also set higher than gains; being set twice (“[2:1]”), five times (“[5:1]”) and ten times (“[10:1]”) higher than gains. The analysis procedure was applied to five clades of reference: Marchantiophyta (i.e., the complete tree), Marchantiopsida, Pelliidae, Jungermanniales, and Porellales. Detailed information on the analyses can be found in Supporting material S1.

**Fig. 3.**
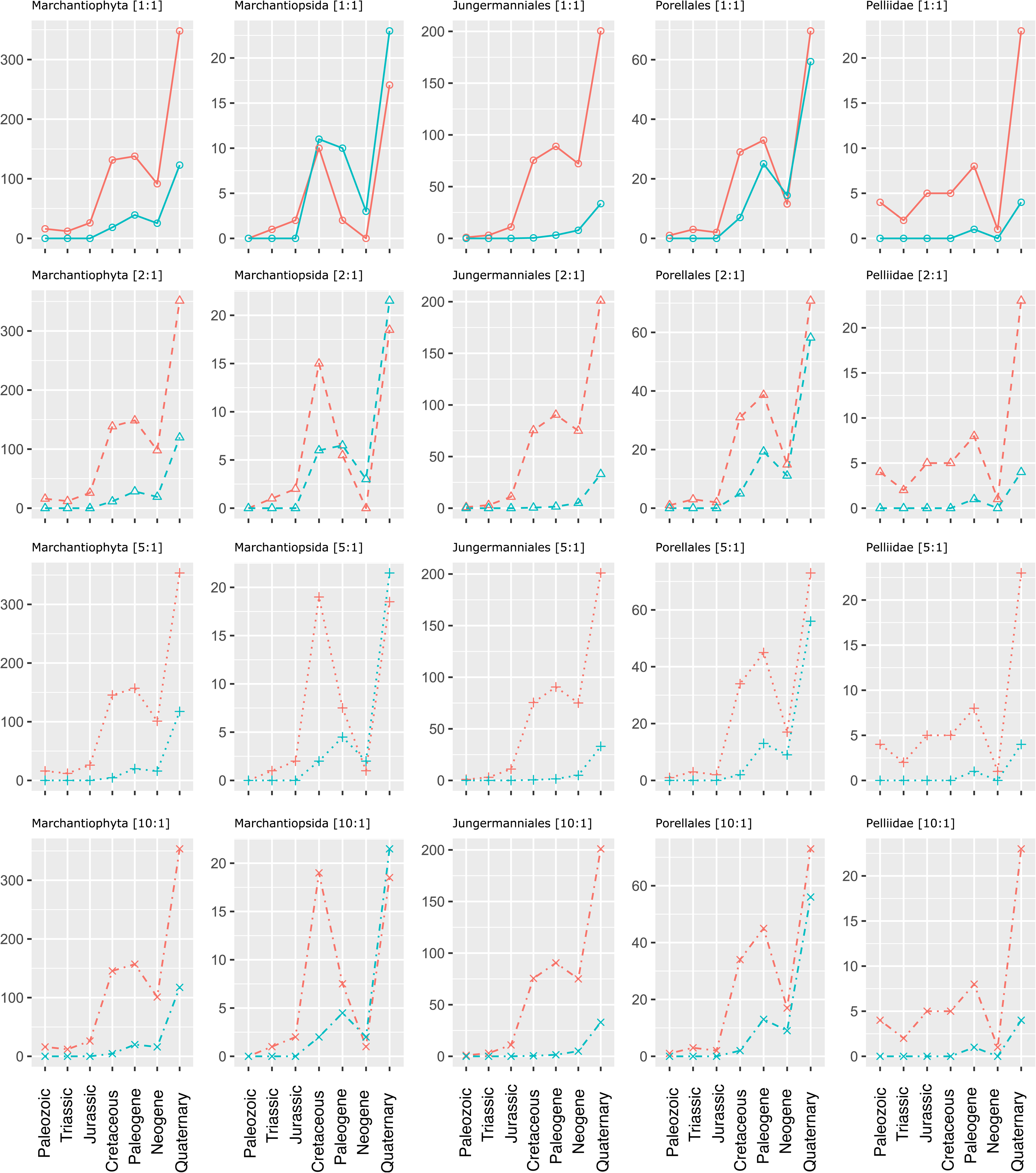
Fluctuation in the evolution of sexual systems in liverworts across geological time. Seven time-bins were defined based on node ages that could be assigned to the following geologic periods: Palaeozoic (> 252 Myr), Triassic (252-201.5 Myr), Jurassic (201.5-145 Myr), Cretaceous (145-66 Myr), Paleogene (66-23 Myr), Neogene (23-3 Myr) and Quaternary (< 3 Myr). The average number of nodes reconstructed as dioicous (orange) and monoicous (blue) are estimated upon inferring ancestral character states onto the MCC dated phylogeny by using maximum parsimony as optimality criterion (Fig. 1). “Reversals” (changes from monoicy to dioicy) were set equal to (“[1:1]”) or higher than “gains” (changes from dioicy to monoicy; “[2:1]”, “[5:1]” and “[10:1]”). The analysis procedure was applied to five clades of reference: Marchantiophyta (i.e., the complete tree), Marchantiopsida, Pelliidae, Jungermanniales, and Porellales. Detailed information on the analyses can be found in Supplementary Material S1. The result derived from “gains” (changes from dioicy to monoicy; “[2:1]”, “[5:1]” and “[10:1]”) are shown in Supplementary Material Fig. S2.

In contrast, most of the shifts from dioicy to monoicy in our analyses occurred in the leafy Porellales (Figs. 1–3) and the complex thalloid Marchantiopsida (Figs. 1–3). In Marchantiopsida, monoicy has appeared after the Jurassic and, depending on the transformation cost, the number of nodes reconstructed as monoicous in this clade has surpassed the dioicous ones already by the Early Mesozoic or Early Cenozoic (Fig. 3). In Porellales, monoicy evolved after the Cretaceous and, when transformation costs are equal between sexual systems, the number of monoicous nodes increased beyond the number of dioicous ones during the Neogene followed by a second shift to dioicy (Fig. 3). Based on these results, our explanation on the dynamic changes in sexual systems is that dioicy is selected in liverworts and that monoicy, despite occurring repeatedly, is less stable than dioicy. Our hypothesis on the evolution of monoicy will be discussed in more detail in a later section.

We show that evolution of sexual systems in liverworts is complicated, being both conserved as seen in the maintenance of dioicy, but also highly dynamic as seen in the shifts of the sexual system. Because dioicy is highly stable and conserved in numerous lineages through time, and in support with available evidence of cytology, we can only assume that the dioicous condition must be linked to the genetically controlled sex-determination. Such sexual systems likely fit most liverworts and all bryophytes having a gametophyte which enables to reproduce asexually and vegetatively in addition to sexual reproduction. It may also partially explain the slow evolution rates found in bryophytes (Linde, 2019) because time for changes between sexual systems is required; that is, a species may need to accumulate some amount of genetic change over time in order to effectively evolve from one sex phenotype to the other. The time lag between the dioicy and monoicy (Fig. 3) supports this hypothesis. If the chromosomal controlled sexual system is highly conserved, knowledge must be acquired through genomic studies on the sex chromosomes in order to assess how dioicy is maintained in the course of evolution.

### The central role of sex chromosomes in bryophyte evolution

Evolutionary implications of sex chromosomes have been extensively studied in diploid organisms, especially in animals (Muller, 1914; Bull, 1983; Rice, 1984; Charlesworth, 1996). Widely accepted model for sex chromosome evolution in diploid organisms with XY or ZW systems suggests that sex chromosomes evolve from a pair of autosomes, initially by acquisition of a sex-determining locus, with subsequent emergence of sexually antagonistic alleles at loci close to the sex-determining locus select for reduced recombination, leading to degeneration of Y/W (Charlesworth, 1996; Bachtrong, 2013). In bryophytes, both empirical and theoretical studies on the evolution of sex-determination and sex chromosomes have been limited. Because of its specific mode of inheritance, the haploid sexual system occurring in bryophytes and also in some macroalgal, as well as fungal systems has been recently designated as UV sex chromosome system (Bachtrog *et al.*, 2011; Coelho *et al*., 2018). In a haploid system, the female U and the male V are either unpaired in the haploid gametophyte and both are exposed to purifying selection, or they are paired in the diploid UV sporophyte. Bull (1978) predicts that the U and V sex chromosomes should show similar characteristics, including similar extent of degeneration, minor degenerations in both, retention of genes on U or V required by the gametophyte and loss of genes required only in the diploid, and that changes in size of U or V should be additions rather than losses. Recent model (Immler & Otto, 2015) predicts that the degeneration is expected to be slower in haploid organisms because the U and V are exposed to haploid selection and also, they do not undergo a marked reduction in the effective population size compared with diploid organisms (one-half for U or V compared to autosomes, but one-quarter for Y/W). However, deleterious mutations in sex-determination genes can be masked if they function in the diploid sporophyte.

Empirical data derived from *Marchantia polymorpha* (Bowman *et al.*, 2017) and *Ceratodon purpureus* (Carey *et al*., 2020) agrees largely with the theories in suppressed recombination on both U and V chromosomes with concomitant repetitive element acquisition. The presence of homologs between the U and V, known as gametologs, also supports the prediction that in a haploid organism the essential genes on sex chromosomes are more likely to persist. *M. polymorpha* has a pair of small, heteromorphic sex chromosomes, including a larger U chromosome (n=8+U) in the female and a smaller V chromosome (n=8+V) in the male, and both sex chromosomes are smaller than the autosomes (Haupt, 1932; Bischler, 1986). Bowman *et al*. (2017) show that *M. polymorpha* exhibits a long-time absence of recombination between the U and V chromosomes pointing to their ancient origin before the split of Marchantiidae and Pelliidae ca. 400 MYA. Likewise, the sex chromosomes have undergone a large degree of degeneration on both sex chromosomes — with a five-fold lower gene density in both U and V than in the autosomes, which does not support Bull’s theory (Bull, 1978).

The *M. polymorpha* 10-Mb V chromosome is characterized by a number of striking features. It was designated into two segments of YR1 and YR2, both are rich in repeats, but the origins of these repeats are very different (Okada *et al*., 2001; Yamato *et al*., 2007). YR1 is composed of copies of repeat sequences consisting of a small number of repeat elements in various arrangements to form an extensive 2-3 Mb V chromosome-specific stretch. A male-specific gene family, ORF162, with an estimation of a few hundred copies, was found embedded in the repeat sequences (Okada *et al.*, 2001; Yamato *et al.*, 2007). This unique feature of co-amplification of protein-coding genes with unique repeat sequences may signify the stage of degeneration of the V chromosome and also indicate how the genes required for male functions have been maintained over the course of evolution (Okada *et al*., 2001; Tanurdzic & Banks, 2004). RY2 of the 6 Mb segment of the V chromosome is composed of repeats and transposable elements accounting for approximately 43% of the segment and is relatively gene rich (Yamato *et al*., 2007). In the most recent analysis, there are 129 annotated genes on the V chromosome, with 19 of these being gametologs shared with the U chromosome (Bowman *et al*., 2017; Montgomery et al 2020). Of the 110 genes unique to the male genome many expressed in reproductive organs but not in vegetative thalli, suggesting their participation in male reproductive functions (Yamato *et al*., 2007; Bowman *et al*., 2017). Six male reproductive protein-coding genes have homologs in animals but not in angiosperms, possibly involved in spermatogenesis as they encode proteins related flagellar components of other species (Okada *et al*., 2001; Yamato *et al.*, 2007; Bowman *et al*., 2017). Many V-specific loci are autosomal genes or gene fragments that have been shown to have recently accumulated into the V chromosome, and the same pattern also occurs for the U, suggesting dynamic evolution of the non-recombining regions of both sex chromosomes (Bowman *et al*., 2017).

The study on chromatin profiling of *M. polymorpha* (Montgomery *et al*., 2020) shows further that the V chromosome is the most densely packed with transposons belonging to all different classes, as suggested in earlier analyses (Yamato *et al*., 2007), and with an abundance of silencing histone modifications, a pattern which is in stark contrast to the relatively uniform interspersion of transposons and genes in autosomes. These observations are molecular confirmation of Heitz’s (1928) cytological analyses nearly a century earlier. The strong compaction may likely have important evolutionary implications in regulating gene and transposon activity relating to sexual differentiation. In animals, there have been ample examples showing that transposable elements can regulate the expression of sexual development genes (Dechaud *et al*., 2019).

As with the V chromosome, the U chromosome is sparsely populated with genes, with only 74 annotated genes, of which 20 have V chromosome gametologs (Bowman *et al*., 2017). There are only a few functionally U-specific genes, including a presumably feminizer locus that can dominantly determine sex in diploid gametophytes (Haupt, 1932; Bowman *et al.*, 2017). Also similar to the V chromosome, the U chromosome harbors specific repetitive sequences in the form of rDNA clusters with distinct intergenic sequences that evolved independently of that on autosomes (Fujisawa *et al*., 2003).

The gametolog pairs shared between the *M. polymorpha* U and V chromosomes exhibit no synteny, despite these genes presumably being descended from genes on the ancestral autosome that gave rise to the sex chromosomes (Bowman *et al*., 2017). Analysis of synonymous substitution frequencies between the members of a gametolog pair can provide a rough estimate of their time of divergence, and thus indirect evidence for sex chromosome evolutionary strata that arises by successive rearrangements incorporating sex chromosome regions into the non-recombining region initially localized to the sex determination locus. Such analyses in *M. polymorpha* indicate multiple evolutionary strata, with the oldest predating the Marchantiopsida-Jungermanniopsida divergence (Bowman *et al*., 2017).

Compared with *M. polymorpha*, the sex chromosomes of *C. purpureus* show structural variation between the U and V to a similar extent, and highly differentiated transposable elements accumulation, but with the non-recombining regions younger (McDaniel *et al*., 2013a; Carey *et al*., 2020). Carey *et al*. (2020) demonstrated that the sex chromosomes of *C. purpureus* expanded via at least two distinct chromosomal fusions to form neo-sex chromosomes and most of the numerous sex-linked genes to the non-recombining U and V are of recent recruitment. The authors suggest that the evolution of sexual dimorphism in bryophytes is largely driven by sexual antagonistic selection through sex chromosome rearrangement, including gene translocations and also sex chromosome translocations in some species (Carey *et al*., 2020). They further showed that in *C. purpureus*, genes involved in sexual development and functions evolved faster than other genes, indicating there is distinct set of evolutionary forces acting on sex chromosomes relative to autosomes. Therefore, sex chromosomes should have profound impact on genome evolution, such as lack of ancient polyploidy in liverworts (Bowman *et al.*, 2017), and likely also on speciation process (Carey *et al*., 2020). This unique feature of sex chromosomes has remained largely unexplored in bryophytes.

While the identity of the U chromosome ‘feminizer’ proposed in early genetics experiments (reviewed in Bowman, 2016; Heitz, 1949) is not yet known, part of the downstream sex-determination pathway of *M. polymorpha* has been elucidated. An autosomal *FEMALE GAMETOPHYTE MYB* (Mp*FGMYB*) was identified as a gene specifically expressed in female plants (Hisanaga *et al*., 2019). Expression of Mp*FGMYB* in males is suppressed by the gene *SUPPRESSOR OF FEMINIZATION* (*SUF*), by producing an antisense RNA at the Mp*FGMYB* locus, thus there is a *cis*□cting bidirectional transcription switch controlling sexual dimorphism. The expression of *SUF* is expected to be suppressed by the unidentified feminizer encoded by the U chromosome (Bowman *et al*., 2017). Whether this sex-determination pathway is *Marchantia* specific, and whether different pathways exist in other liverwort species, more species should be investigated. The autosomal *cis*-acting sexual dimorphism switch Mp*FGMYB* was found to be orthologous to the recently captured U and V-linked Mp*FGMYB* gene copies in the moss *Ceratodon purpureas* (Carey *et al*., 2020). Therefore, the genomic evidence suggests that the sex chromosomes of *M. polymorpha* is relatively preserved, and their evolution is highly dynamic. These studies mentioned above provide genetic evidence for sex-specific genes on the sex chromosomes, that is, there likely exist multiple ‘motility’ loci on the V encoding male-specific proteins required for flagellar function and that at least one locus on the U for egg cell development.

Theoretical evidence suggests that if sex-antagonistic genes are located on autosomes, sexual antagonistic mutations will be selected to be linked to the sex-specific nonrecombining region through chromosomal rearrangements with autosomes (Charlesworth & Charlesworth, 1980; Rice, 1984). Bull’s theory on the sex chromosome in haploid dioicy (Bull, 1978) was proposed mostly based on the cytology and genetics of liverwort genus *Sphaerocarpos* possessing a chromosome set n = 8 (7+U/V) (Knapp, 1936; Allen, 1945). His prediction, however, on the equal magnitude of degeneration of the sex chromosomes did not fit the karyotypes of the genus in which the U chromosome is larger than both the autosomes and V chromosome, and the V is smaller than the autosomes. Bull assumed that this discrepancy suggests that there is some fundamental difference between the male and female gametophyte or their gametes which favors these additions in the female but not the male chromosome. However, this is unlikely as the opposite conditions also exist.

*Sphaerocarpos* is in many ways different from *Marchantia*. It occurs widely in warm and dry Mediterranean climate and has an ephemeral habit to avoid extreme conditions by having a short-lived gametophyte, and with a sporophyte producing large and resistant spores that lie dormant in the driest months (Schuster, 1992). Some *Sphaerocarpos* species shed their spores in tetrads, hence keeping two males and two females together — thus, even though it lives in an ephemeral habitat, it has not evolved monoicy, but rather evolved another mechanism to ensure that males and females are growing together.

The life history of *Sphaerocarpos* implies that much of the resource of the female should be allocated to the sporophyte development, thus suggesting parent-offspring conflict. In his experimental studies on the inheritance of gametophytes of *Sphaerocarpos*, Allen (1919, 1935) reported that none of the mutant genes he found on the gametophytes was borne on a sex chromosome but there was a puzzling amount of linkage of certain mutants with sexuality, therefore, he assumed that certain autosomes tend to be associated with sex chromosomes during meiosis. Although cytological evidence shows that the U chromosome is much larger and more heterochromatic than the V, Meyer and Herrmann (1973) demonstrated using reassociation analysis that about 22% of the DNA of *S. donellii* is repetitive and there is no difference in the repetition of nucleotide sequences between DNA in males and females. Furthermore, in the attempt of identifying sex specific markers in *S. texanus*, surprisingly few markers (three as specific to females and one to males) were found (McLetchie & Collins, 2001). These early findings suggest that the sex chromosomes of *Sphaerocarpos* underwent large scale rearrangements, likely through sex chromosome–autosome fusion to form neo-sex chromosomes. It has been shown that in animal species larger and more heteromorphic sex chromosomes are associated with faster evolution of postzygotic isolation, leading to divergence, thus can contribute to ecological specialization and speciation (Paladino *et al*., 2019). Therefore, it seems clear that genetically controlled sex-determination in bryophytes is maintained through a highly dynamic evolutionary process, and as in other sexual systems predicted by van Doorn & Kirkpatrick (2007), that accumulation of sex-antagonistic polymorphisms may enhance evolutionary stability of the long-established sex-determination system and at the same time it may also promote speciation.

### Reversal to dioicy, less stable monoicy

Sexual differentiation in monoicous species depends on their immediate environment, and there is no or little constraint on spatial isolation of the sexes, therefore, monoicy must have adaptive advantages, such as enhanced possibility for sexual reproduction, and hence, dispersal. It has been shown that intra-gametophytic selfing, which occurs frequently in monoicous bryophytes, can efficiently prevent the accumulation of deleterious mutations (Szövényi *et al*., 2014). On the other hand, repeated events of intra-gametophytic selfing may hinder adaptive evolution as predicted by theory (Birky & Walsh 1988; Charlesworth, 2012).

As we have shown in the previous section, shifts of sexual system from dioicy to monoicy occurred mainly in recently diverged nodes within Marchantiopsida and Porellales clades, wherein monoicy persisted over time in many genera (Figs. 1–3). Laenen *et al.* (2016) showed that monoicous lineages have higher diversification rates than dioicous lineages in liverworts, stating that increased diversification rate follows the shift to monoicy. Although species of Marchantiopsida and Porellales have a different evolutionary history, distribution and physiology, they share habitats that are often unstable or temporary wherein species should have higher growth rates and be able to complete their life cycles soon after favorable conditions are set. Therefore, monoicy is an adaptation of liverworts to such habitat. Species of both lineages tend to exhibit a higher level of parental investment on sporophyte by producing larger spores, for surviving during unfavorable intervals. Species of Porellales are mostly epiphytes growing on tree trunks and/or living leaf surfaces in humid tropical forests. Their spores undergo precocious germination within the capsule, with their release as a several-celled, chlorophyllose sporeling ready for immediate further development (Schuster, 1983). In the study of sexual system evolution of genus *Radula*, Devos *et al*. (2011) showed that transitions to monoicy from the dioicous ancestral condition were phylogenetically significantly correlated with epiphytism and that it is not the sexual system that determines the evolution of epiphytism, but the reverse. In many species of Marchantiopsida that occur in seasonally dry areas, their gametophytes have either evolved short life cycles and with sporophytes producing durable spores, or both the gametophytes and sporophytes become drought resistant linked with morphological adaptations to reduce water loss, for example, the development of ventral parenchymatous tissue and dorsal assimilatory chlorenchyma of the thallus, and their spores with thick and rigid walls in addition to the large size (e.g., in *Mannia*, Schuster, 1992). Therefore, monoicous expression is likely driven by certain life histories.

There has been no ancient whole genome duplication (WGD) retained in liverworts (Bowman *et al.*, 2017), but so far it is known that one molecular mechanism leading to monoicy is through a polyploidization event resulting in both U and V sex chromosomes being present (Berrie, 1964; Ramsay and Berrie, 1982). However, because some monoicous liverworts appear to be haploid based on cytology (Berrie, 1960) it is likely that monoicy can evolve through sex chromosome rearrangement, for example, by the U chromosome feminizer translocating to an autosome. In monoicous species, genes responsible for controlling sex expression may be dispersed throughout the genome. Note that a monoicous species derived from ancestral dioicious species might retain a chromosome, now autosomal, that descended from a sex chromosome but that has characteristics of an H or h chromosome. Comparative studies on both dioicous and monoicous species, especially closely related species pairs would likely provide insights into the genomic changes associated with transitions between sexual systems.

Although monoicy has evolved repeatedly in liverworts, reversal to dioicy did occur in some species of both Marchantiopsida and Porellales. The cause for the resultant pattern is not manifest, but because the time interval for the transition to dioicy from monoicy is short (Fig. 3), we presume that monoicy is less stable in the long-term than dioicy in liverworts. This may also be the case in mosses and hornworts, as it has been shown that in mosses the transition rate from monoicy to dioicy was approximately twice as high as the reverse transition (McDaniel *et al*., 2013b). To test this hypothesis, the impact of sexual systems of bryophytes on island biogeography can provide some hints. Patiño *et al.* (2013) found that the proportion of monoicous taxa was significantly higher on islands, and that a significant proportion of continental species that are monoicous or dioicous are represented on oceanic islands only by monoicous populations. They further pointed out that the life history traits shifted toward a greater proportion of species producing asexual propagules and smaller proportion of species producing spores, showing weakened advantage of monoicy over time. In other studies, reduced fitness among progeny produced by selfing in monoicous mosses under certain stressful conditions have been suggested (Jesson *et al*., 2012). Hornworts show an extreme paucity in species diversity with little over 200 species in total, and except for species of the genus *Dendroceros* that have evolved as epiphytes growing on tree trunks and their gametophytes can tolerate drier periods, the rest species lack desiccation tolerance and many of them grow on moist soil in transient habitats as annuals (Wood, 2007; Warny *et al.*, 2012). It is thus not surprising that there are more monoicous species than dioicous ones. Furthermore, Villarreal and Renner (2013) found that the transition rate from dioicy to monoicy was twice higher than in the opposite direction, but monoicous groups have higher extinction rates.

Unlike previously assumed that monoicous species should have larger ranges than dioicous species (Longton & Schuster, 1983), Laenen *et al.* (2015) found that sexual systems are not correlated with geographical ranges, and they suggested that monoicous species can also experience severe fertilization constraints under certain conditions. A better understanding on the evolutionary significance of monoicy in liverworts is still necessary. At present, we could argue that the “erratic” or “constant” shifts in monoicy through time (Fig. 3) are reasonable based on Patiño *et al.*’s and Jesson *et al.*’s findings, and they are further supported by Laenen *et al.*’s suggestion. Since monoicy poses constraints on fertilization (Laenen *et al*., 2015) and offers “weak” advantages over time (Patiño *et al*., 2013; Jesson *et al*., 2012), the tendency of the nodes being monoicous fluctuates through evolutionary time, thus, dioicy is ultimately favoured (Fig. 3).

### Directions for future research

A number of bryophytes have long served as model plants in studies of cytology, cytogenetics and genetics, which led to the remarkable findings of plant sex chromosomes and heterochromatin. It was once predicted by the well-known botanist and geneticist of his time, Fritz von Wettstein (1895-1945), that bryophytes would “remain especially favored organisms for many geneticists” (Wettstein, 1932). Unfortunately, Wettstein’s prediction did not turn to be true until the present day mainly because of the lack of sufficient study tools and techniques in the early times. Genetic basis for phenotypic evolution can now be addressed using genome-wide information to further advance our understanding of reproductive biology of bryophytes, which is of vital importance in studies of evolution, biodiversity, systematics, development, ecology, as well as conservation, among many others. Both genetical factors and environmental influence play an important role in the evolution of sexual systems, the implication of the former has been little assessed for bryophytes, therefore, the following points can be put forth for the future studies.

First, more genome characterization among key bryophyte species is to be accumulated for allowing comparative studies of the evolution of sexual systems including sex-determination, sexual dimorphism, and testing existing hypotheses and also hypotheses proposed herein, in order to build up evolutionary basis on sexual reproduction. It is expected that sex chromosomes should play a large role in sexual dimorphism, thus can further shape their evolutionary and genomic properties. In a broader scale, UV sexual system can provide substantial contribution to the understanding of the evolution of sex chromosomes. For bryophytes, there is still a lack of knowledge regarding why one sexual system is more favored over the other, and particularly, knowledge about how the reversal of sexuality occurs, i.e., how a monoicous species with autosomes descended from sex chromosomes can evolve back to dioicy and whether this is easier than evolving sex chromosomes de novo or, alternatively, relatively equivalent. These gaps may be filled with further investigations on relevant groups such as families Frullaniaceae, Radulaceae and Ricciaceae.

Second, cytological and functional differences between sex chromosomes should be more studied. Because the evolution of the sex chromosomes in bryophytes is much more dynamic than thought before, cytological and cytogenetical studies may likely provide new information on the potential of sex chromosome rearrangement and turnover. Phenotypic evolution of dioicy and monoicy in combination with habitat conditions, and life history traits may be further studied in light of the increased knowledge of nature and evolutionary significance of the sex chromosomes. This will lead to a deeper understanding on the spatial and temporal distribution of the bryological diversity. In the field of taxonomy and systematics, R. M. Schuster realized long ago that taxonomical problems such as species delimitation is expected to be solved with increased knowledge of reproductive biology and biogeography (Schuster, 1988).

Third, the above proposed endeavor will also help understand the evolution of polyploidy in bryophytes, thus the species diversity. So far, genetic implications of polyploidization event on the evolution of sex chromosomes, such as whether the rarer occurrence of polyploidy in liverworts than in mosses is related to the stability of dioicy, and to the female dominated expression in the diploid gametophytic phase, wait further studies.

Fourth, sexuality of a species should be considered as a key feature if it is subjected for conservation among other life history traits. Dioicous and monoicous species may be susceptible to threats in different way and to different extent, thus, these factors should be considered in the conservation effort. Conservation strategies will become meaningless if information in reproductive biology of the species to be protected is inadequate, as reproduction together with dispersal is the key element affecting whether a species will be sufficiently resilient to climate change or become vulnerable to extinction.

## Acknowledgements

JF acknowledges support by FONCyT (grant PICT 0810), and PIUNT (grant G631).

## Notes

### Competing Interest Statement

The authors have declared no competing interest.

### Summary of Updates

1.Only a few typo errors (one in Abstract, and in figure captions) were corrected. 2. three missing references were added to the reference list. One was deleted. 3. Author name John Bowman changed to John L. Bowman 4. A few typo errors were corrected, and one sentence was added in the supporting material S1.

