## Supplemental Materials and Methods S1 for "Living apart if you can – how genetically and developmentally controlled sex has shaped the evolution of liverworts"

Dated Phylogeny reconstruction

The phylogenetic relationships across liverworts were reconstructed by using 466 species in 299 genera and 84 families (out of 376 genera and 87 families accepted worldwide, Söderström *et al*., 2016) based on Bayesian Inference analysis, as implemented in BEAST v1.10.4 (Suchard *et al*., 2018). Thirteen species of mosses were used as the outgroup. Of 3232 aligned bases of nucleotides represent three markers (*rbc*L, *rps*4 and *trn*L-F). The sequence data were downloaded from the GenBank.

Divergence time estimation was inferred in a Bayesian framework using BEAST v1.10.4 (Suchard *et al*., 2018). To estimate absolute ages for lineage divergences we used seven fossil points based on fossil records within liverworts to set minimum age constraints for several nodes in the phylogeny. These fossils include Marchantites cyathodoides (225 Myr, Townrow, 1959), Cheirorhiza brittae (158 myr, Krassilov, 1970), Diettertia montanensis (112 Myr, Brown & Robison, 1974), Nipponolejeunea europaea (50 Myr, Grolle, 1981), Mastigolejeunea contorta (50 Myr, Grolle & Meister, 2004), Calypogeia stenzeliana (50 Myr, Grolle & Meister, 2004), and Bazzania polyodus (50 Myr, Grolle & Meister, 2004)*.* According to the hypothesis that liverworts arose close to the earliest fossil record of land plants (Kenrick and Crane, 1997; Wellman *et al*., 2003), we applied a normal distribution prior with a mean of 475 Mya and an SD of 5 Mya, which contains 95% of the probability distribution between 465 and 485 Mya. Minimum age constraints were enforced at each calibration points by applying an exponential prior from the fossil age to 475 Mya (the maximum age of the root of Marchantiophyta). We used the uncorrelated lognormal relaxed clock model to prevent the negative effects from heterogeneity of substitution rates and uncertainty of fossil data (Drummond *et al*., 2006). GTR+Γ+I was used as substitution model with Gamma Categories set to 6, and a Yule tree prior was used as the tree model. MCMC chains were run for 100 million generations with parameters and trees logged every 10000 generations. Tracer v1.5 (Rambaut and Drummond, 2007) was used to check the parameters and the trees were combined in TreeAnnotator v1.10.4 with 1% burnin using the value 100. The final single tree visualized using FigTree v1.4.4 (Figure S1).

An alternative method using previously published plastid substitution rate was also conducted with the smaller dataset to cross-validate the inferred divergence times by using fossils. According to the studies of Palmer (1991) and Schnabel and Wendel (1998) who report average absolute rates of substitutions in cpDNA of 5.0 10– 4 substitutions− 1⋅site − 1⋅Myr across a wide range of algae and land plants, we applied a normal distribution with a mean of 5.0 10– 4 and standard deviation of 10− 4 substitutions− 1⋅site − 1⋅Myr as a prior on the absolute *rbc*L, *rps*4 and *trn*L-F rates of evolution. The analysis was done with BEAST v1.10.4 (Suchard et al., 2018)).

Ancestral character state reconstruction

In this study, the evolution of the sexual system through time was examined by mapping a binary discrete character (dioicous (0), monoicous (1)). Information of the sexuality of the sampled species was obtained from literature. Although the Mk model (Lewis, 2001) has gained attraction to conduct phylogenetic analyses based on morphological data or combined data types (i.e., “total evidence approach”), it has been recently shown that the model assumptions are hardly met by morphology (Goloboff and Arias, 2019; Goloboff *et al.*, 2019); thus, questioning its utility to analyze such data type. Therefore, instead of relying on model-based approaches, we inferred ancestral character states (“ACSs”) across time bins by using parsimony as optimality criterion. Ancestral character states were reconstructed onto the MCMC tree from the divergence time estimation using parsimony as implemented in the program MESQUITE 3.6.1 (Maddison and Maddison, 2019).

In order to evaluate the evolutionary dynamics of the sexual system in liverworts, ACS were reconstructed in TNT 1.5 (Goloboff and Catalano, 2016). In TNT, the ACSs for the sexual system, as defined above, were obtained by optimising the binary character onto the MCMC tree derived from the divergence time estimation analysis. “Reversals” (changes from monoicy to dioicy) were first set equal to “gains” (from dioicy to monoicy; “[1:1]”). However, to take into account possible scenarios, reversal costs were set higher than gains; being set twice (“[2:1]”), five times (“[5:1]”) and ten times (“[10:1]”) higher than gains. Since nodes can be assigned ambiguous ACSs (i.e., different character states can be equally optimal), we employed a 1000-round iterative approach wherein ambiguously optimized nodes were solved at random. In each cycle of this procedure, the number of nodes optimized as *dioicous* or *monoicous* were summed up and averaged for each of the discretely-defined time bins. This procedure was applied to five clades of reference: Marchantiophyta (i.e., the complete tree), Marchantiopsida, Pelliidae, Jungermanniales, and Porellales.

Seven time-bins were defined based on node ages that could be assigned to the following geologic Era or Periods: Palaeozoic (> 252 Myr), Triassic (252-201.5 Myr), Jurassic (201.5-145 Myr), Cretaceous (145-66 Myr), Paleogene (66-23 Myr), Neogene (23-3 Myr) and Quaternary (< 3 Myr). Because of the few nodes that were older than 252 Myr, we used a wider time-bin (Palaeozoic) for such nodes. Conversely, since the number of younger nodes was higher, we employed “thinner” time-bins (i.e., Periods) to increase the resolution of the assessment through time. To define time-bins in TNT, first, node ages were sequentially estimated as the difference between branch lengths and the age of nodes ancestor; starting from the root node age as implied by the MCC phylogeny. Subsequently, nodes were assigned to the respective time-bins according to the distribution of their ages across time. The whole procedure (i.e., node ages estimation, assignation of nodes to time-bins and ACS reconstruction per time-bin and clade) was implemented in a TNT script available upon request.
