## Supplementary figures and images for "Living apart if you can – how genetically and developmentally controlled sex has shaped the evolution of liverworts"

### Supplemental Figure S1

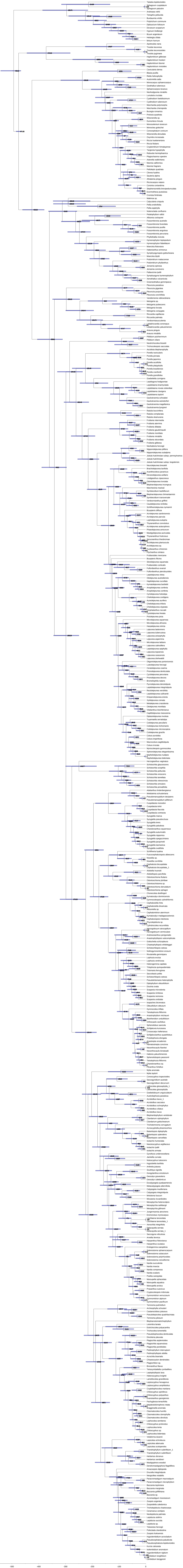

### Supplemental Figure S2

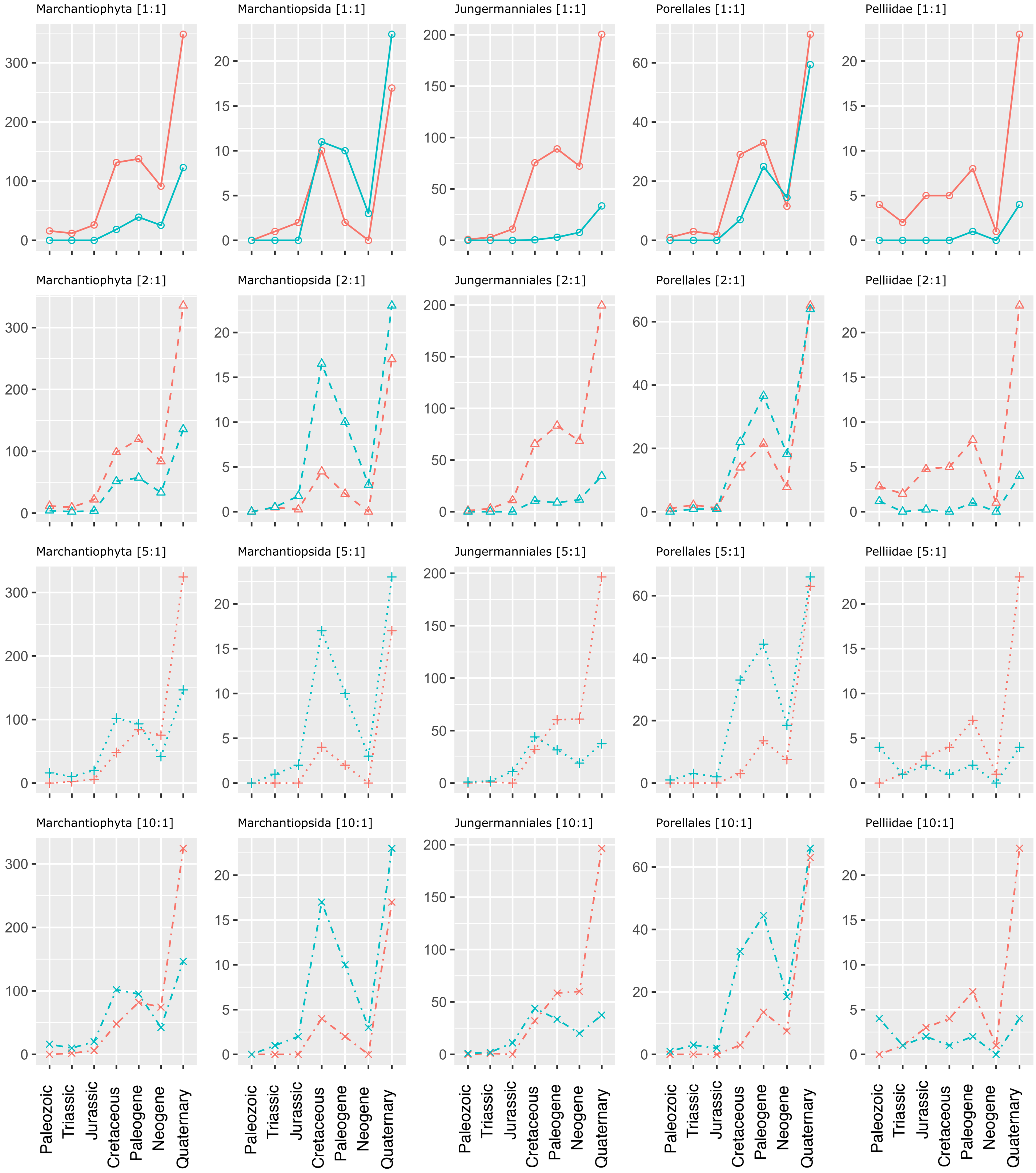
